# Quantitative imaging reveals PI3Kδ inhibition reduces rhinovirus-induced damage of small airway epithelia in ex vivo cultured human precision cut lung slices from COPD patients

**DOI:** 10.1101/2022.03.01.482451

**Authors:** Dmytro Dvornikov, Aliaksandr Halavatyi, Muzamil Majid Khan, Natalie Zimmermann, Amy Cross, Daniel Poeckel, Emil Melnikov, Christian Tischer, Johannes Leyrer, Marc A Schneider, Thomas Muley, Hauke Winter, Edith Hessel, Soren Beinke, Rainer Pepperkok

**Affiliations:** Cell Biology and Biophysics Unit, European Molecular Biology Laboratory, 69117 Heidelberg, Germany; Cellzome GmbH, GSK, 69117 Heidelberg, Germany; Translational Lung Research Center Heidelberg (TLRC), German Center for Lung Research (DZL), 69120 Heidelberg, Germany; Advanced light microscopy facility, European Molecular Biology Laboratory, 69117 Heidelberg, Germany; Medicines Research Centre, GSK, Stevenage SG1 2NY, UK; Faculty of Biosciences, Heidelberg University, 69120 Heidelberg, Germany; Translational Research Unit, Thoraxklinik at University Hospital Heidelberg, 69126 Heidelberg, Germany; Department of Surgery, Thoraxklinik at University Hospital Heidelberg, 69126 Heidelberg, Germany

## Abstract

Chronic obstructive pulmonary disease (COPD) is one of the major causes of disability and death worldwide and a significant risk factor for respiratory infections. Rhinoviral infections are the most common trigger of COPD exacerbations which lead to a worsening of disease symptoms, decline in lung function and increased mortality. The lack of suitable disease models to study the relevant cellular and molecular mechanism hinders the discovery of novel medicines that prevent disease progression in exacerbating COPD patients. We used quantitative multi-color imaging of COPD and control patient derived human precision-cut lung slices (hPCLS) to study the impact of rhinovirus infection on the structure and function of the small airway epithelium. Data analysis highlighted that COPD-derived hPCLS have a higher cellular density and basal cell hyperplasia, more unciliated airway surface areas with mucus overproduction, and shorter cilia length compared to control hPCLS. In response to rhinovirus 16 infection, COPD-derived hPCLS secreted higher amounts of pro-inflammatory cytokines and displayed decreased epithelial integrity and reduced airway ciliation. Finally, treatment with a selective PI3Kδ inhibitor reduced secretion of rhinovirus-induced cytokines and ameliorated rhinovirus-induced damage to COPD small airway epithelia. Thus, these data demonstrate the potential of quantitative imaging to assess complex airway functions in a patient-derived lung tissue model system, and indicate that targeting PI3Kδ might be a promising therapeutic opportunity to limit rhinovirus-induced airway damage in exacerbating COPD patients.

**Summary:** PI3Kδ inhibition reduces rhinovirus-mediated damage of small airway epithelia from chronic obstructive pulmonary disease (COPD) patients

## INTRODUCTION

Chronic obstructive pulmonary disease (COPD) is the third leading cause of death worldwide. The incidence and mortality rates of COPD are rising due to environmental factors like smoking and air pollution^1^. The disease is characterized by progressive and irreversible airflow obstruction, chronic airway inflammation and destruction of the lung parenchyma. COPD patients can be highly susceptible to common viral and bacterial respiratory infections which trigger exacerbations of disease symptoms and drive disease progression, with each subsequent episode further worsening the lung function, thus increasing morbidity and mortality^2^. Rhinovirus infections are the main cause of virus-triggered exacerbations in COPD patients^3,4^. Triple therapy, inhaled corticosteroid (ICS), a long-acting β2-agonist (LABA), and a long-acting muscarinic antagonist (LAMA) in a single inhaler is the current standard of care to manage the severity and frequency of acute exacerbations^5^. However, novel therapeutics that prevent the exacerbation associated decline in lung function is needed to improve patient outcomes.

The airway epithelium is the first line of defense that provides a biophysical barrier against pathogens and noxious agents^6^. Multiple epithelial cell subsets make up the pseudo-stratified epithelium that in small airways mainly consist of basal, ciliated and secretory cells^7^. The airway epithelium is the primary site of rhinovirus infection in the lungs^8^ and infected airway epithelial cells activate interferon responses and inflammatory pathways that attract and activate immune cells^9^. Increased lung inflammation and mucus production is a hallmark of COPD exacerbations. While altered barrier and cilia function has been observed in COPD patients^10^ this has not been extensively studied in the context of HRV infection.

Phosphoinositide 3-kinase delta (PI3Kδ) is a lipid kinase that is preferentially expressed in leukocytes where it regulates cell activation, differentiation and proliferation, thereby modulating immune responses^11^. PI3Kδ expression and activation is elevated in neutrophils and lung tissue from COPD patients and its level of expression correlates with the disease severity^12^. Neutrophils isolated from COPD patents demonstrate increased migration speed alongside with reduced directionality toward the cytokine gradient which was reversible by PI3Kdelta inhibition^13,14^. Importantly, patients with PI3Kδ activating mutation (APDS syndrome) develop recurrent respiratory infections and chronic lung inflammation^15,16^. In line with this, conditional knock-in mice with a PI3Kδ activating mutation display higher susceptibility to bacterial infection, while PI3Kδ inhibition reduces pro-inflammatory cytokine secretion and improves survival rates following infection^17^. These lines of evidence make PI3Kδ an attractive therapeutic target for COPD.

Lack of a suitable COPD model system has hindered development of novel therapeutics, e.g. the establishment of relevant clinical endpoints^18^. Use of primary cells and cell lines is usually limited to single cell types or simple co-cultures grown on two-dimensional substrates, while mouse models reflect only certain aspects of the disease due to mouse-to-human differences in lung physiology, immune system composition and impact of environmental and life style factors^19,20^. The precision-cut lung slice (hPCLS) model has recently emerged as a powerful tool to study complex human diseases as multiple cell types are preserved in their natural three-dimensional microenvironment that allows in-deep investigation of complex physiological tissue responses^21,22^. Here, we aimed to directly compare the responses of small airways and surrounding parenchyma to rhinovirus infection in COPD and control patient derived hPCLS. Furthermore, using combined multiplexed cytokine release assays and quantitative multi-color imaging we investigated whether PI3Kδ inhibition modulates the response to rhinovirus infection. The results demonstrated that treatment with a PI3Kδ inhibitor significantly reduced the elevated secretion of pro-inflammatory cytokines and small airway epithelium damage in rhinovirus-infected COPD hPCLS.

## RESULTS

### Quantitative multi-colour imaging of human airways and automated analysis

We first aimed to setup an imaging workflow that enables comprehensive multi-parametric quantitative analysis of human small airways in large numbers. We have previously shown that hPCLS are morphologically heterogenous^22^. To fully capture heterogeneity of human small airways we embedded 250-µm thick airway-containing hPCLS between two coverslips to allow two-sided confocal imaging (fig. 1a). Low magnification imaging of hPCLS derived from COPD patients before *ex vivo* culture demonstrated classical hallmarks of the disease: marked destruction of lung parenchyma (fig. 1b) alongside with collapsing small airways containing mucus plugs (fig. 1c). To assess the phenotypic differences in airway cellular composition, we performed immunostaining against ADP Ribosylation Factor Like GTPase 13B (ARL13B) for labelling cilia, Tumor Protein P63 (P63) as a marker of basal cells andMucin 5AC, Oligomeric Mucus/Gel-Forming (MUC5AC) for labelling mucus and mucus producing cells. High magnification imaging of the small airways allowed more thorough comparison of COPD and non-COPD small airway epithelium structure. Non-COPD small airways contained pseudostratified columnar epithelium with almost equally spaced basal cells positioned on the basal lamina. In contrast, COPD small airway epithelium showed disorganized epithelium with hyperplasia of airway epithelial cells and frequent clusters of basal cells (white asterisk, fig. 1d). Additionally, COPD epithelium contained more mucus-producing cells and had overall higher levels of mucus (white arrowheads, fig. 1d). We performed 3D reconstruction of the airway surface to allow better visual comparison of the ciliation differences between COPD and non-COPD small airways (Black asterisk fig. 1e). This analysis revealed higher prevalence of non-ciliated regions in COPD epithelium.

**Figure 1.**
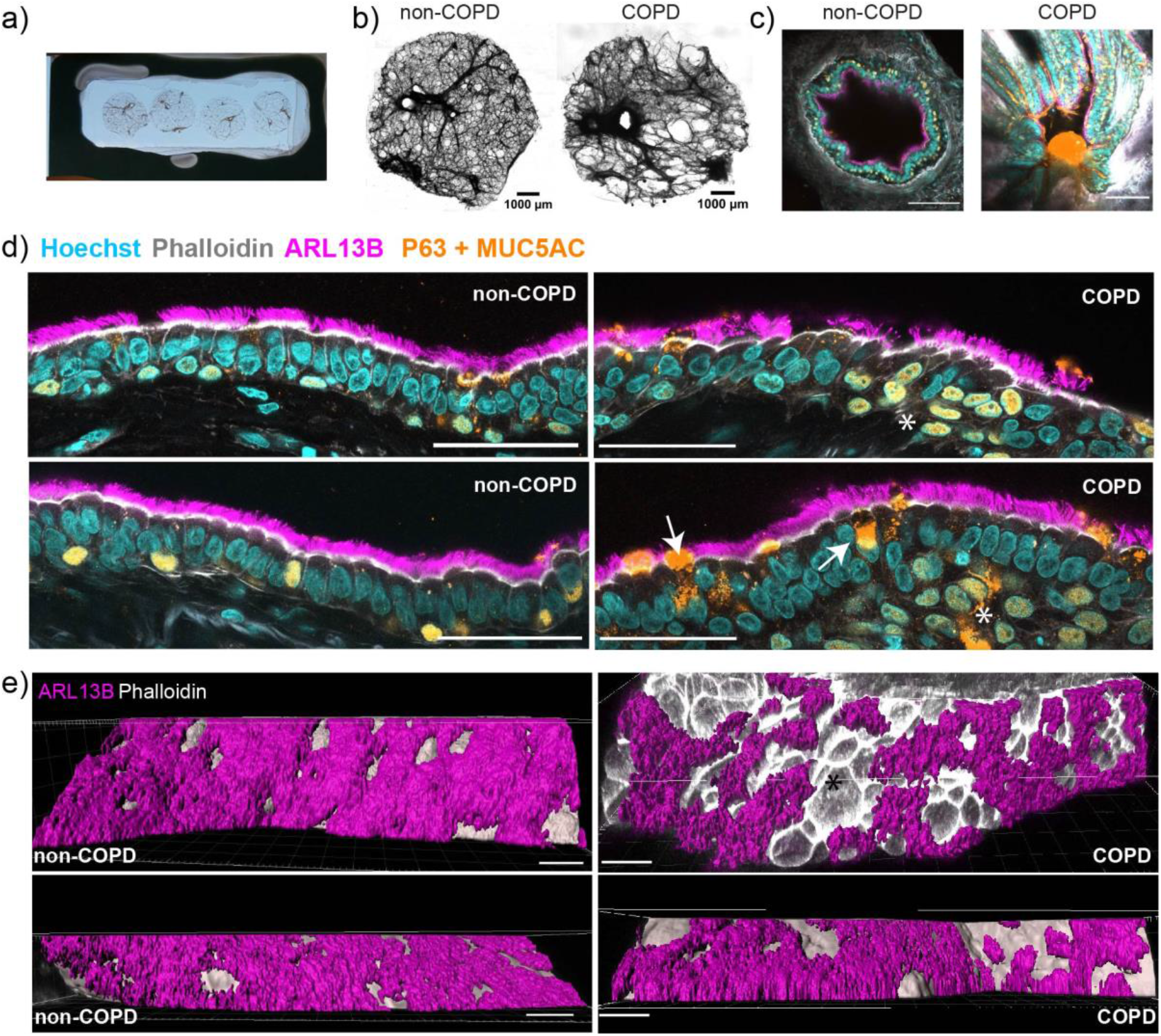
Quantitative confocal multi-color imaging of whole-mount human precision-cut lung slices (hPCLS). a) Small airway containing hPCLS embedded between to coverslips to allow two-sided confocal imaging. b) Representative images of airway hPCLS from non-COPD and COPD donors. c) Representative images of non-COPD and COPD small airways. Colors the same as in d. Scale bar 100 µm. d) Representative confocal images of non-COPD and COPD small airway epithelium. Scale bar 50 µm. e) 3D reconstruction of non-COPD and COPD small airway epithelium. Scale bar 20 µm. Color schemes are for the same protein markers in c & d. White arrow head= Mucus, white asterisk= clusters of basal cells, black asterisk= non-ciliated regions

We then aimed for unbiased quantitative comparison of small airway epithelium state across multiple z-stacks, airways and patients. We established a Java-based automated image analysis pipeline that is implemented as a stand-alone ImageJ plugin. The pipeline allows quantification of multiple quantitative parameters: i) percentage of ciliated airway surface; ii) number of total cells covering epithelial surface; iii) number of basal cells; iv) number of cells in the airway lumen; v) total cilia area and intensity; vi) total mucus area (fig. 2a). Importantly, results of automated and manual quantification of several parameters including percentage of ciliated airway surface, total number of epithelial cells and number of basal cells were highly correlated (R^2^ = 0.87–0.98) (fig. 2b). We also assessed whether automated analysis of individual 2D images that belong to the same z-stack results in a similar outcome as the manual analysis of the same 3D reconstructed z-stack. Indeed, percentage of ciliated airway surface and the percentage of the basal cells were highly comparable between both approaches (fig. 2c). Therefore, in all subsequent analyses of confocal airway images 2D automated quantification was used as it offers higher throughput without compromising the data quality.

**Figure 2.**
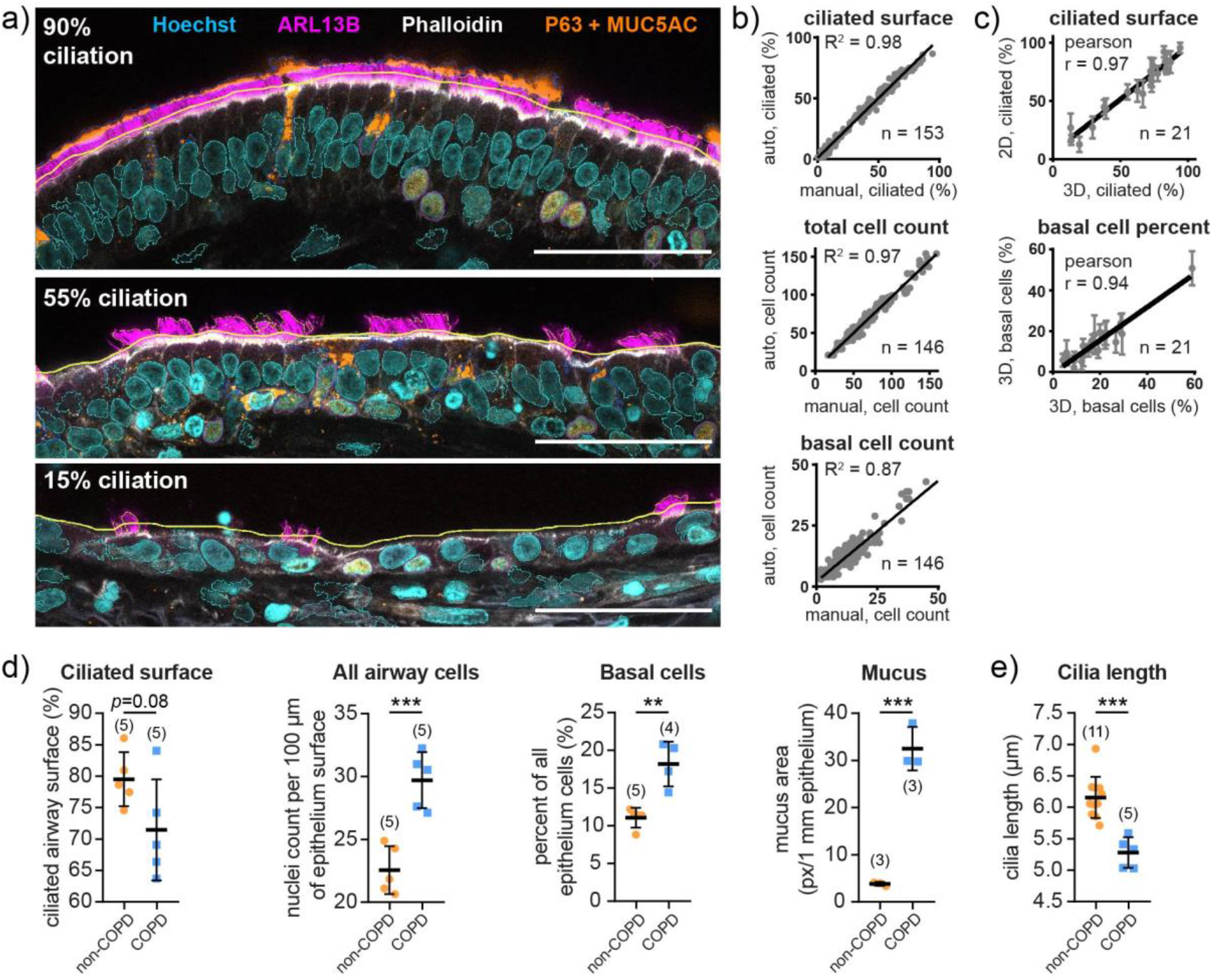
Pipeline for automated analysis of confocal images. a) Representative images with different degrees of ciliation analyzed with the pipeline. Thick yellow line depicts the airway border automatically selected based on Phalloidin staining. Non-basal cell nuclei are shown with cyan contour, basal cells with magenta contour, mucus with blue contour, cilia regions with thin yellow contour. Number indicates percent of ciliated airway surface as measured by the pipeline. Scale bar 50 µm. b) Linear regression of manual and automatic analysis results of the ciliated airway surface, number of total and basal cells. n is the number of images that were used for comparison. c) Linear regression of 2D and 3D analysis results of the ciliated airway surface and percent of basal cell population. For 2D analysis mean and SD was calculated for each z-stack. n is the number of z-stacks that were used for comparison. d-e) Comparison of percentage of cililated airway surface, total number of cells in the airway epithelium, percentage of basal cells, mucus area and cilia length in untreated airway non-COPD and COPD hPCLS using automated (d) and manual analysis (e). Number in the brackets corresponds to the number of donors per group. For each donor median of all quantified images across several airways is shown. Full distribution of each quantified parameter, number of airways and z-stacks used for analysis for each patient is presented in supplementary figure S1. Data presented as mean and SD. Statistical analysis by unpaired *t*-test. ***: *P*<0.001; **: *P*<0.01.

We applied the established automated image analysis pipeline to quantitatively compare COPD and control airways from baseline hPCLS. Combined analysis was performed on a total of five COPD patients (29 airways, 321 z-stacks, 68.2 cm total airway surface, ∼208,000 epithelium cells) and five control patients (28 airways, 241 z-stacks, 88 cm total airway surface, ∼194,000 epithelium cells). The airway surface of COPD patients had less ciliation (69.0 ± 8.0% vs. 79.5 ± 4.3%, *P* = 0.08) than that of control patients (fig. 2d and supplementary figure S1b). The COPD small airway epithelium contained significantly more cells per 100 um of surface length (30.5 ± 2.2% vs. 21.8 ± 1.9%, *P* = 0.0006), a higher percentage of basal cells (18.8 ± 3.0% vs. 11.4 ± 1.3%, *P* = 0.0018) and higher levels of mucus (299 ± 46 px/mm vs. 40 ± 4 px/mm, *P* = 0.0004) (fig. 2d and supplementary figure S1c-e). Additionally, manual analysis of cilia length between five COPD patients (∼44,500 individual cilia across 23 airways) and eleven control patients (∼41,400 individual cilia across 39 airways) and showed significantly shorter cilia in COPD airways (5.3 ± 0.24 µm vs. 6.2 ± 0.32 µm, *P* = 0.001) (fig. 2e and supplementary figure S1a).

### Human precision-cut lung slices (hPCLS) are viable with functional small airway upon *ex vivo* culture

Next, we aimed to optimize hPCLS culture conditions to ensure high hPCLS viability, to facilitate HRV infection and PI3Kδ inhibitor treatment while minimizing the presence of additional ambient stimuli that can potentially confound our observations. In order to assess whether hPCLS required foetal calf serum (FCS) as a culture medium component, hPCLS were cultured for three days either in the presence of 5% FCS, 1% FCS, or none. FCS addition did not affect mitochondrial activity of hPCLS as determined by WST-1 assay (fig. 3a), and therefore was omitted from all subsequent experiments. We then assessed time-dependent changes in hPCLS viability by measuring lactate dehydrogenase (LDH) leakage into the culture medium over a three day-time period (fig. 3b). Amount of released LDH each day remained constant and below 10%, with hPCLS derived from two COPD donors showing higher LDH release than hPCLS derived from non-COPD donors (fig. 3c). Next, we aimed to investigate whether our established *ex vivo* culture conditions affect cilia beating frequency (CBF) in small airways present in hPCLS (fig. 3d). We established fully automated Python-based CBF analysis pipeline that allows unbiased batch analysis of CBF recordings (fig. 3e and f). There was no significant difference (paired *t*-test, *P* = 0.10) in CBF measured using the established pipeline and manual counting (fig. 3g). Given ciliary beating frequency has been shown to be temperature dependent^23^, and increase with increasing temperatures, next we tested the correlation between CBF and changes in temperatures (fig. 3h). Indeed, the pipeline was also able to quantitatively capture the temperature-dependent changes in CBF measured in the same airways (fig. 3h), thus demonstrating that CBF in airway hPCLS responds in an expected manner to temperature perturbations. Finally, we used the pipeline to show that CBF did not change (paired *t*-test, *P* = 0.83) during three days in culture of airway-containing hPCLS derived from four patients (fig. 3i).

**Figure 3.**
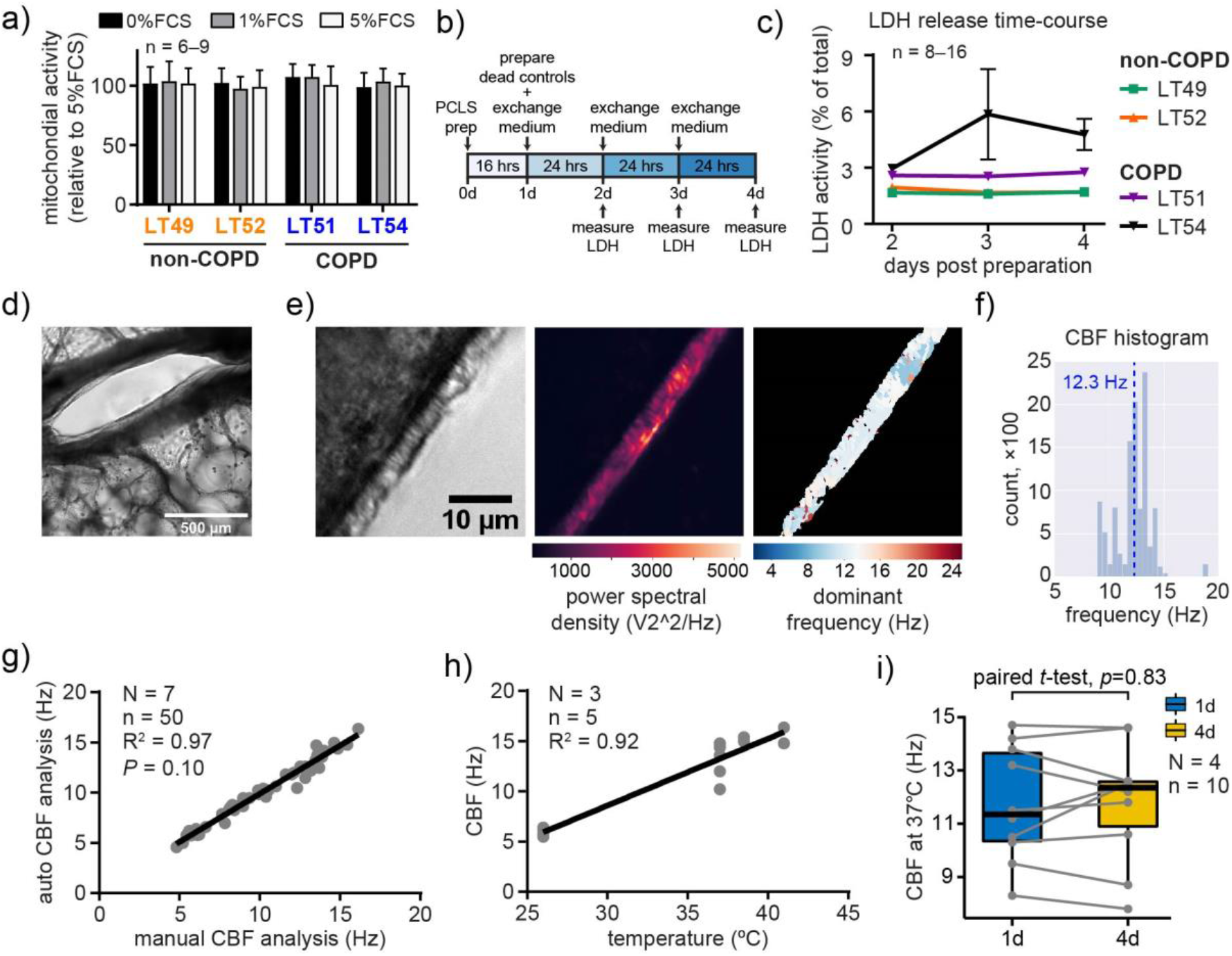
hPCLS maintain high viability and ciliated epithelium function up to four days in culture. a) Evaluation of hPCLS mitochondrial activity using WST-1 viability assay. hPCLS from two COPD and two non-COPD donors were cultured in varying FCS concentrations for 3 days after an initial overnight recovery. Data correspond to mean and SD, n is the number of hPCLS used per condition per donor. b) Experimental setup for assessing hPCLS viability in a time-dependent manner. c) Time-dependent LDH release assessed in two COPD and two non-COPD donors. Medium was exchanged every 24 hrs. Results are presented as a percentage of total LDH as calculated from calibration curve prepared from Saponin-extracted hPCLS. Data correspond to mean and SEM, n is the number of hPCLS used per condition per donor. d) Representative low-magnification image of the small airway. e) Left, representative high-magnification single frame from the high-speed imaging used for automated cilia beating frequency (CBF) analysis. Middle, fast Fourier transform (FFT) power spectrum of an entire image. Right, CBF of pixels with the power above the threshold is color-coded. f) Representative frequency histogram from one automatically-analyzed CBF movie. Dashed blue line shows the median of the distribution. g) Linear regression of the median CBF measured across all video recordings and all airways of a given patient using an automated CBF analysis pipeline compared with manual counting beating events using kymograph analysis. N is the total number of donors, n is the total number of hPCLS. Statistical analysis by paired *t*-test. h) Linear regression of the median CBF measured from the same hPCLS at different temperatures. N is the total number of donors, n is the total number of hPCLS. i) Change in median CBF after 3 days in culture measured from the same hPCLS. Center lines show the medians; box limits indicate the 25th and 75th percentiles. N is the total number of donors, n is the total number of hPCLS.

### Rhinovirus infection induces stronger pro-inflammatory response and cytotoxicity in COPD hPCLS

We established a viable hPCLS culture system with functional small airways over four days in culture, which retained COPD-derived hPCLS disease-specific alterations. We then aimed to compare response of COPD and control hPCLS to increasing doses of rhinovirus to identify the optimal infection dose. hPCLS from two COPD and three control patients were infected with human rhinovirus 16 (RV16). hPCLS from both COPD and control donors showed a dose-dependent increase in cytotoxicity when assessed by the release of LDH (fig. 4a). Importantly, COPD-derived hPCLS released 2–3 times more LDH when exposed to the same virus dose and had overall stronger cytotoxic responses than control hPCLS (*P* < 0.0001). Next, to test the pro-inflammatory does response of hPCLS derived from non-COPD and COPD patients to RV16 infection, we measured the levels of key antiviral cytokines^24^ including interferon gamma-induced protein 10 (IP-10), interferon gamma (IFNγ), interleukin (IL)-1β, IL-2, IL-8, IL-10, macrophage inflammatory proteins 1-alpha and 1-beta (MIP-1α and MIP-1β), and thymus and activation regulated chemokine (TARC) (fig. 4b–f and supplementary figure S2). While both non-COPD and COPD patient derived hPCLS showed a dose-dependent release of key antiviral cytokines (fig. 4b–f and supplementary figure S2), importantly, virus-infected COPD hPCLS consistently released higher levels of pro-inflammatory cytokines.

**Figure 4.**
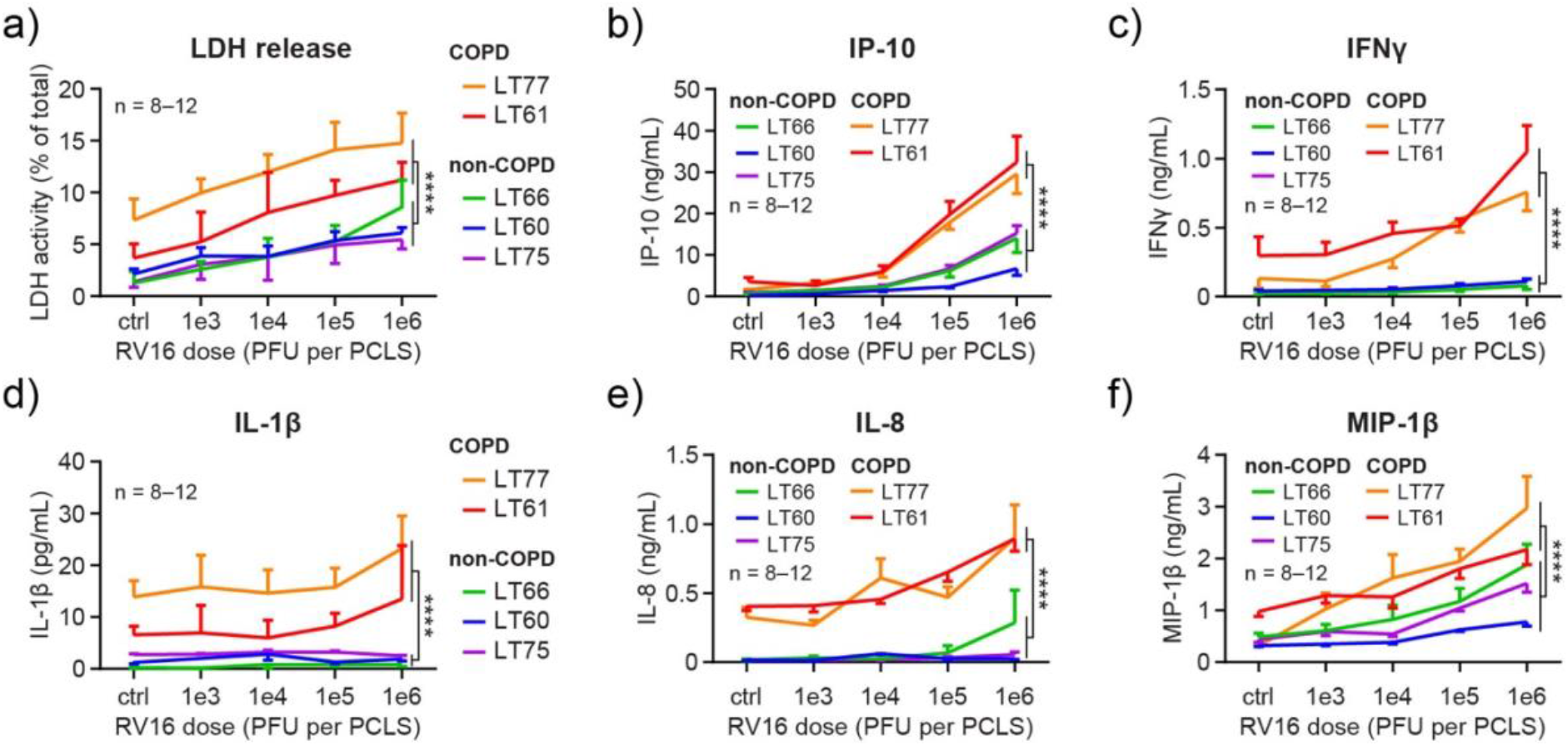
Rhinovirus infection induces higher dose-dependent cytotoxicity and pro-inflammatory response in COPD-derived hPCLS. hPCLS from three non-COPD and two COPD GOLD IV donors were infected with indicated amounts of rhinovirus 16 (RV16). Supernatants were collected 3 days after infection and analyzed for a) LDH activity and b-f) cytokine levels. Data correspond to mean and SEM, n is the number of hPCLS used per dose per donor. Statistical analysis by Factorial ANOVA with Bonferroni multiple comparison *post-hoc* test. *; **; ***; **** indicate significance with *P* < 0.05; 0.01, 0.001; 0.0001. Full results of statistical analyses are shown in supplementary table S2.

### PI3Kδ inhibition attenuates rhinovirus-induced damage of human COPD small airway epithelium

We then aimed to investigate how pre-treatment with a highly selective PI3Kδ inhibitor^25^ would impact the rhinovirus infection. We decided to use the infection dose of 1e5 plaque-forming units (PFU) per hPCLS as it was resulting in pronounced antiviral response, while still preserving COPD airway epithelium in a state acceptable for microscopy analysis. In contrast, a dose of 1e6 PFU resulted in almost complete airway epithelium shedding in COPD hPCLS (supplementary figure S3). Importantly, the PI3Kδ inhibitor alone had no cytotoxic effect in all tested concentrations (fig. S4a and b) and did not impair CBF and bead transport velocity (fig. S4c–e). We therefore treated hPCLS with the highest tested dose of the PI3Kδ inhibitor, 100 nM, followed five hours later by infection with previously optimized RV16 dose of 1e5 PFU per hPCLS. We profiled the responses of hPCLS from two COPD donors (44 airways, 414 z-stacks, 64.5 cm total airway surface, ∼157,700 epithelium cells) and three control donors (62 airways, 618 z-stacks, 100 cm total airway surface, ∼250,000 epithelium cells) and (fig. 5 and supplementary figure S5). PI3Kδ inhibition alone did not cause any significant change in airway ciliation and in the number of cells within the airway epithelium or in its lumen in either control or COPD airways (fig. 6a–c and Supplementary table S3). Three-day-long RV16 infection resulted in a drop of airway ciliation from 79% to 41% in COPD airways (P < 0.0001) and from 82% to 62% in control airways (*P* < 0.0001) (fig. 6a and Supplementary table S3). Importantly, combining PI3Kδ inhibition with RV16 infection significantly reduced loss of ciliation from 41% to 53% in COPD airways (*P* < 0.0001, fig. 6a, Supplementary table S3), while it had no impact on loss of ciliation in control airways (*P* = 0.91). Additionally, in control airways RV16 infection resulted in only a minor drop in number of epithelium cells per 100 µm of the airway surface (from 24.4 to 19.7, *P* < 0.0001, fig. 6b) that was also accompanied by a small increase in a number of cells present in the airway lumen (from 0 to 0.25 cells per 100 µm airway, *P* < 0.0001, fig. 6c and (Supplementary table S3). In contrast, in COPD airways RV16 caused pronounced reduction in airway epithelium cell number (from 30.7 to 14.9 cells/100 µm, P < 0.0001, fig. 6b) alongside a strong increase in number of cells present in the airway lumen (from 0 to 0.9 cells/100 µm, *P* < 0.0001) (fig. 6c and Supplementary table S3). In turn, co-treatment with the PI3Kδ inhibitor reduced RV16-induced loss of epithelium cells from 19.7 to 23.9 cells/100 µm in control airways (*P* < 0.0001), and from 14.9 to 19.2 cells/100 µm in COPD airways (*P* < 0.0001, fig. 6b). Additionally, PI3Kδ inhibition also reduced infection-mediated increase in cells in the airway lumen in both COPD airways (from 0.9 to 0.6 cells/100 µm, *P* < 0.0001) and control (from 0.25 to 0.04 cells/100 µm, *P* < 0.05, fig. 6c). Importantly, pan-hPCLS cytotoxicity assessed by LDH release did not show significant effect of PI3Kδ inhibition on reduction of rhinovirus-mediated cytotoxicity (Supplementary figure S6), thus strongly highlighting the power of our established microscopy-based analysis pipeline.

**Figure 5.**
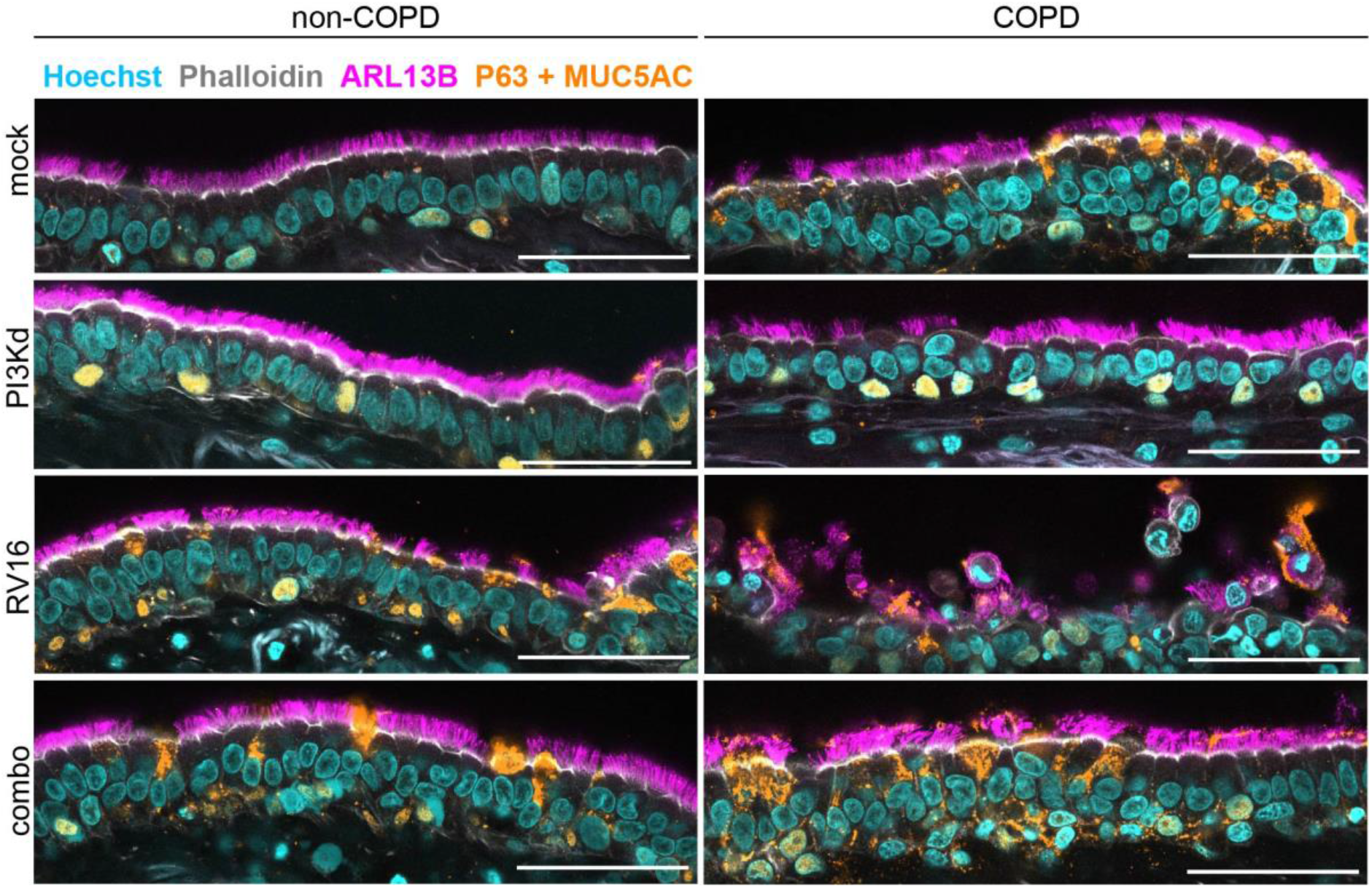
PI3Kδ inhibition differentially impacts integrity of COPD small airway epithelium after rhinovirus infection. hPCLS from three non-COPD and two COPD GOLD IV donors were left untreated (mock), or treated with 100 nM PI3Kδ inhibitor (PI3Kd), or infected with 1e5 PFU/hPCLS of rhinovirus 16 (RV16), or first pre-treated with PI3Kδ for five hours and then infected with RV16 (combo). Three days post infection hPCLS were PFA-fixed, stained with corresponding antibodies and imaged on a confocal microscope. Representative images for each condition of non-COPD and COPD airways are shown. Additional images are presented in supplementary figure S5. Scale bar 50 µm.

**Figure 6.**
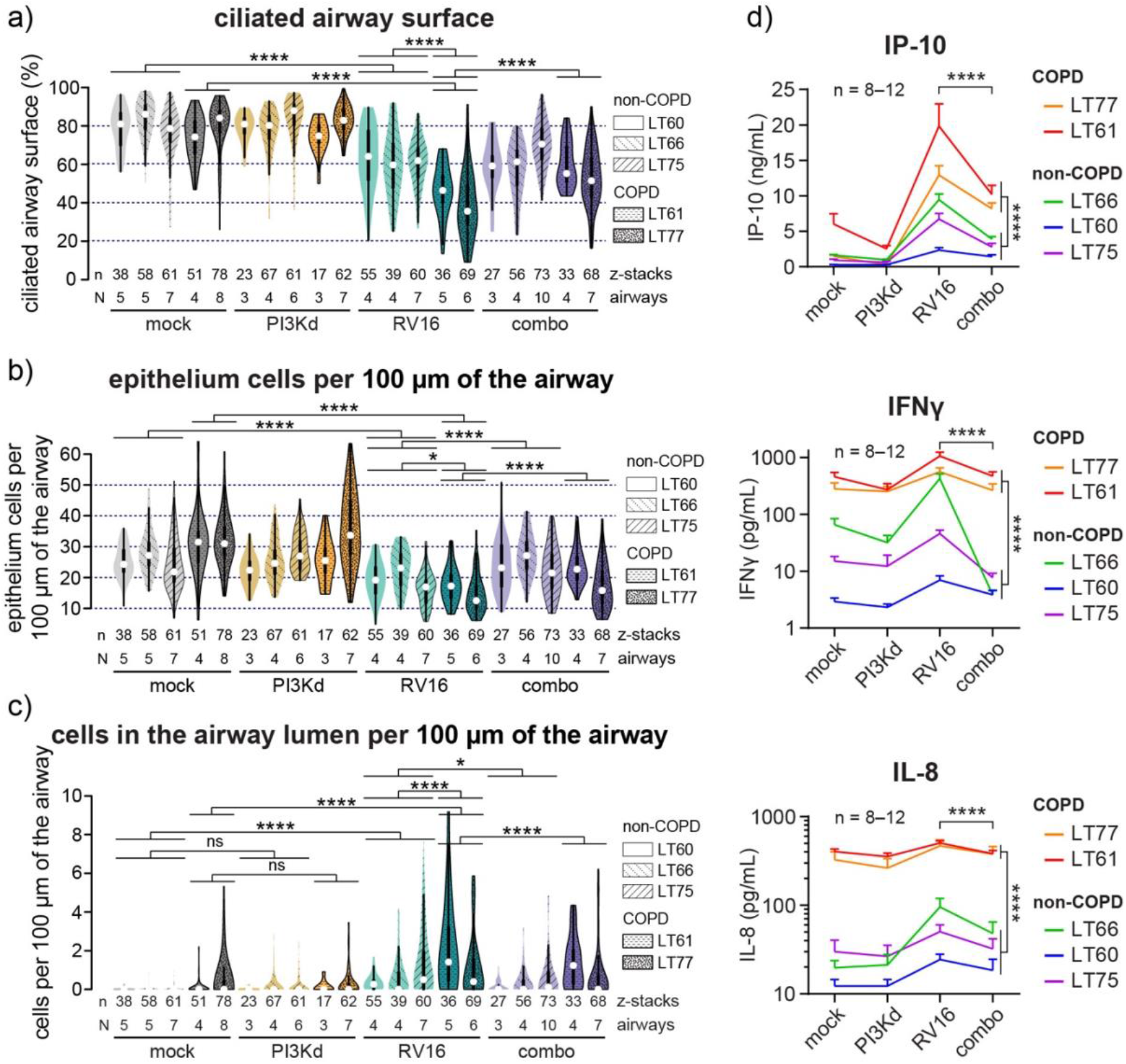
PI3Kδ inhibition attenuates rhinovirus-induced damage of human COPD small airway epithelium. hPCLS from three non-COPD and two COPD GOLD IV donors were left untreated (mock), treated with 100 nM PI3Kδ inhibitor (PI3Kd), infected with 1e5 PFU/hPCLS of rhinovirus 16 (RV16), or pre-treated with PI3Kδ for five hours and then infected with RV16 (combo). Three days post treatment hPCLS were PFA-fixed, stained with corresponding antibodies and imaged on a confocal microscope. a-c) Quantification of percent of ciliated airway surface (a), airway epithelium cell number (b), and number of cells in the airway lumen (c) in non-COPD and COPD hPCLS. White circles show the medians; box limits indicate the 25th and 75th percentiles; whiskers extend 1.5× the interquartile range from the 25th and 75th percentiles; polygons represent density estimates of data and extend to extreme values. d) Cytokine levels measured from hPCLS supernatants 3 days after treatment. Data correspond to mean and SEM, n is the number of hPCLS used per dose per donor. Statistical analysis by Factorial ANOVA with Bonferroni multiple comparison *post-hoc* test. *; **; ***; **** indicate significance with *P* < 0.05; 0.01, 0.001; 0.0001. Full results of statistical analyses are shown in supplementary tables S3 and S4.

We also assessed how PI3Kδ inhibition affects rhinovirus-induced cytokine secretion profile. The PI3Kδ inhibitor did not significantly change the baseline secretion levels of IP-10, IFNγ and IL-8 in non-infected COPD and control hPCLS (fig. 6d and Supplementary table S4). However, when PI3Kδ inhibition was combined with RV16 infection, the induced secretion of all three cytokines was reduced two-fold in comparison to RV16-infected hPCLS. Importantly, hPCLS derived from COPD patients had significantly higher baseline levels of IP-10, IFNγ and IL-8 and also demonstrated stronger induction in response to infection.

## DISCUSSION

Here, we present a quantitative imaging platform that allows unbiased analysis of *ex vivo* treated human airway human precision-cut lung slices (hPCLS). Using the established pipeline, we quantitatively profiled more than a hundred small airways from 16 different COPD patients or control donors. We show for the first time, that selective PI3Kδ inhibition attenuates rhinovirus-induced damage of the human small airway epithelium. Hence, this work showcases the power of quantitative imaging-based hPCLS analysis for dissecting mechanisms of complex human diseases and for clinical translation of promising therapeutic compounds. The results of this work suggest that PI3Kδ inhibition might be a vital option for reducing severity of COPD exacerbation, and thus slowing the disease progression and extending patients’ life expectancy.

In our study, for the first time, we present direct dose-dependent comparison of rhinovirus-infected hPCLS derived from COPD and control primary human lung tissue. Very few studies that characterize responses to rhinovirus infection in COPD patients have been conducted. Experimental rhinovirus infection was performed on healthy human volunteers to assess clinical symptoms and inflammatory mediators in sputum and bronchoalveolar lavage, but more comprehensive molecular analysis was not possible and COPD patients were excluded^4,26,27^. Other studies performed in vitro infection on monocultures of isolated primary bronchial epithelial cells or immune cells from healthy or COPD donors^14,28–30^. Therefore, our work bridges the gap between both approaches by using *ex vivo* cultured hPCLS that retain multiple lung resident cell types that bear COPD-specific alterations and allow detailed mechanistic studies. We show that the profile of released pro-inflammatory chemo- and cytokines (including IP-10, IFNγ, IL-1β, IL-8 and MIP-1β) by COPD hPCLS in response to rhinovirus infection is similar to control hPCLS, but substantially amplified. These findings are in line with several previous studies that have used in vitro and animal models, and human subjects to show that release of IP-10 and IL-8 chemokines was consistently upregulated in COPD context^14,26,28,29,31,32^. Importantly, our results shed light on controversies regarding other cytokines released by COPD lung tissue. Here, we show that rhinovirus infection results in higher IL-1β secretion from COPD-derived hPCLS. In line with our findings, elevated IL-1β was reported in sputum and serum of exacerbating COPD patients compared to healthy smoking and non-smoking controls^4,27^, although primary epithelial cells isolated from COPD patients did not show upregulation of IL-1β in response to rhinovirus infection^28^. Additionally, we demonstrate that both IL-10 and IL-13 are secreted in significantly larger amounts by virus-infected COPD hPCLS. Previous studies report conflicting results on both cytokines. COPD mouse models showed lower IL-10 and elevated IL-13 levels following virus infection^31,33,34^, while COPD patients with naturally occurring virus infection had higher IL-10 and unchanged IL-13 serum levels than control subjects^35^. In our work, we show that rhinovirus-infected COPD hPCLS release higher levels of IFNγ. Previous studies show highly divergent results. In line with our observation, virus infection of *ex vivo* human lung tissue resulted in higher level of IFNγ secretion in COPD^30^. In contrast, virus-infected primary epithelial cells isolated from COPD and control donors released comparable IFNγ levels^28^, while mouse COPD model reported unchanged^36^, increased^31,34^ or even decreased IFNγ^33^.

Our study is the first to report that selective PI3Kδ inhibition ameliorates rhinovirus-induced damage to COPD airway epithelium. We also demonstrate that RV16 infection inflicts limited damage to control airway epithelium that is in line with several previous studies^37–39^. We show that pre-treatment with PI3Kδ inhibitor reduced infection-mediated loss of airway epithelial cells and airway ciliation, and reduced secretion of pro-inflammatory chemo- and cytokines in both COPD and control hPCLS. Previous reports have shown that application of a PI3Kδ inhibitor improved survival rates and decreased IFNγ, IL-1β and IL-10 levels following S. pneumoniae infection in mice with an activating PI3Kδ mutation that symptomatically resembles the COPD phenotype^17^. Similarly, non-selective PI3K inhibition has been demonstrated to reduce IP-10 and IL-8 secretion by RV16-infected primary bronchial epithelial cells and IL-8 secretion by LPS-stimulated COPD blood neutrophils^40^, while selective PI3Kδ inhibition reduced LPS- and TNFα-induced secretion of IL-8 by monocytes from patients with COPD^12,41^, and attenuated IFNγ and IL-2 cytokine secretion by in vitro co-cultures of T- and B-cells from healthy donors^42^. Furthermore, administration of an inhaled PI3Kδ inhibitor to COPD patients with stable disease reduced sputum inflammatory mediators such as IL-8 and IL-13^43–45^ and a clinically meaningful improvement in FEV1^45^. However, the same PI3Kδ inhibitor-induced effects were less pronounced in COPD patients experiencing an acute exacerbation^45^. We further extend these findings by showing for the first time that PI3Kδ inhibition prior to RV16 infection results in reduction of human airway epithelium damage alongside with decreased secretion of several pro-inflammatory cytokines. We hypothesize that in future studies our data and work flow pipeline in hPCLS together with development of highly potent and selective inhibitors of PI3Kδ^25^ will further add to the clinical relevance of PI3Kδ inhibition in prevention of COPD exacerbations.

We performed direct quantitative comparisons of the small airway epithelium composition across more than 140 small airways from five COPD and eleven control patients with a combined analyzed airway surface exceeding 300 cm. We show that COPD small airway epithelium in average contains 40% more total cells (30.5 vs 21.8 cells/100 µm), 65% higher percentage of basal cells (18.8% vs. 11.4%) and shorter airway cilia (6.0 vs. 5.3 µm). In line with our data, several previous studies have similarly shown that smokers and COPD subjects have shorter cilia length^46–48^. Although such studies often measured only around a hundred of individual cilia from about a dozen of ciliated cells per patient. In our work, we measured the length of more than 85,000 individual cilia across 16 patients to show that COPD patients have reproducibly lower cilia length. Furthermore, our measurements of the number of total epithelial cells and basal cells is in line with the work of Boers et al. who systematically assessed cell composition of non-COPD human airways (22.3 cells/100 µm and 10%, respectively)^49^. While it is widely accepted that the COPD airway epithelium features pronounced abnormalities, including such COPD hallmarks as basal cell hyperplasia and reduced number of ciliated cells^50^, comprehensive quantitative assessment of these features remained limited. Here, our automated image analysis pipeline will provide the required quantitative assessment.

COPD is a complex disease that involves deregulated cross-talk of multiple immune and epithelial cell types^51^. Such complexity is difficult to reproduce using conventional in vitro models that hinders discovery of relevant disease mechanisms and assessing the impact of novel therapeutic approaches. Therefore, one of the main strengths of our work is the use of a patient-derived *ex vivo* hPCLS model system that retains multiple disease-relevant cell types with imprinted disease states. COPD is also a notoriously heterogeneous disease with pronounced intra- and inter-patient variability. Different small airways of the same lung and even different regions of the same airway can be differentially affected^50^. Therefore, another important strength of our work is comprehensive imaging and analysis of the airways (an average of 13.4 airways, 1000 images covering 22 cm of airway surface with 57,000 epithelium cells per patient) that allowed reliable unbiased automated quantitative profiling of disease alteration and treatment-induced changes in the small airways. While our experimental approach does address thoroughly intra-individual heterogeneity of COPD patients, patient access was the main limitation to include more COPD samples in the current study. However, our work may allow imaging to be included as an endpoint in future clinical research studies.

PI3Kδ controls neutrophil activation^52,53^, regulates NADPH-dependent superoxide production by monocytes^54^, and production of pro-inflammatory cytokines^55^. A selective inhibition of these over-activated immune cells in COPD lungs is likely to reduce the damage to lung epithelial tissue inflicted by excessive proteolytic activity of the deregulated COPD immune cells. Therefore, in our study, we hypothesize that the mechanism of reduced rhinovirus-mediated airway epithelium damage following PI3Kδ inhibition is highly likely due to the reduction of collateral damage by the activated immune cells. COPD lung resides in a persistent pro-inflammatory state with increased infiltration of macrophages, neutrophils, mast cells and T-cells^56,57^. COPD-derived neutrophils and macrophages reside in the pre-activated state and secrete increased levels of pro-inflammatory mediators and proteolytic enzymes, mast cells release activators of pro-matrix metalloproteinases^57–59^, while increased secretion of pro-inflammatory cytokines by T-cells further drives excessive inflammatory responses^30^. However, the exact identity and contributions of the immune cell types affected by the PI3Kδ inhibition in *ex vivo* human hPCLS still remains to be elucidated. Additionally, it would be important to understand whether distinct COPD endophenotypes differentially respond to PI3Kδ inhibition.

Altogether, our study demonstrates that PI3Kδ inhibition in human COPD hPCLS suppresses rhinovirus-mediated secretion of pro-inflammatory chemo- and cytokines followed by reduced damage of small airway epithelium. These observations might have important clinical implications in developing COPD-specific therapeutic intervention strategies to ameliorate severity of virus-induced exacerbations in COPD patients, thus improving patients’ quality of life and slowing progression of the disease. Our work also showcases the advantages of using hPCLS model system in combination with quantitative imaging in mechanistic studies of complex human diseases in living tissue *ex vivo*.

## MATERIALS AND METHODS

### Human lung tissue

The experiments with human tissue were approved by Ethics Committee at Medical Faculty of Heidelberg University (reference number S-270/2001) and by EMBL Bioethics Internal Advisory Committee (references BIAC 2017-001 and 2018-004). The human biological samples were sourced ethically and their research use was in accord with the terms of the informed consents under an IRB/EC approved protocol. Written informed consent was obtained from all subjects. Tumor-free lung tissue was obtained from early-stage lung cancer patients who underwent lung lobe resection with curative intent. Tissue was received and prepared within one hour after surgery. Clinical characteristics of the patients are provided in the Supplementary Table S1.

### Preparation of human hPCLS

Lung tissue (4–20 g) was inflated with warm low temperature gelling agarose (3% by weight in DMEM w/o Phenol red, A9414, Sigma) at room temperature using 1-mL syringe. Tissue was kept for 30 min in ice-cold DMEM to allow gelling of the agarose. Cylindrical 8-mm diameter tissue cores were prepared using semi-automated coring tool (Alabama Research and Development, USA). Cores were trimmed on both sides to remove damaged tissue after the syringe-inflation. 250-μm thick hPCLS were produced using Krumdieck tissue slicer (Alabama Research and Development, USA) filled with ice-cold DMEM w/o Phenol red. Immediately after slicing, hPCLS were washed twice in warm DMEM-BSA (w/o Phenol red, 100 U/ml penicillin, 100 μg/ml streptomycin, 2 mM L-glutamine, 1 mg/mL BSA and 2.50 µg/mL Amphotericin B) for 30–60 min in the incubator (37°C, 5% CO2, 100% humidity). Following the washing steps hPCLS were transferred into 24-well plates (one hPCLS per well) and kept overnight in 1 mL DMEM-FCS (same composition as DMEM-BSA but with 5% FCS instead of BSA) in the incubator. The following day medium was replaced with 1 mL of DMEM-BSA.

### Rhinovirus infection and PI3Kδ inhibitor treatment

hPCLS were infected for one hour in 100 µL MEM (Gibco) in 24-well plates at 37°C with human rhinovirus 16 (Virapur, kindly provided by GSK RRI DPU, Stevenage, UK). Infection dose ranged from 10^3^ to 10^6^ PFU per hPCLS depending on a condition. PI3Kδ inhibitor (GSK2269045C) was applied five hours prior to infection, and was refreshed after infection. DMSO was used as a vehicle control. After infection hPCLS were washed twice with DMEM-BSA, and incubated for three days in 1 mL DMEM-BSA per hPCLS in the incubator (37°C, 5% CO2, 100% humidity).

### Cytotoxicity assays

hPCLS viability was assessed by measuring lactate dehydrogenase (LDH) activity and using WST-1 cell proliferation kit. LDH activity in the supernatants was measured using CytoTox 96 kit (Promega, G1780) according to the manufacturer’s instructions. 50 µL of supernatant per slice was used. To convert relative LDH activity values to absolute values (% of total LDH released) total LDH content of the representative hPCLS was extracted by incubating slices in DMEM-BSA with 0.1% Saponin at 37°C for one hour. Resulting supernatants were used to make five-point dilution curve for converting relative activity values to absolute LDH content values. For WST-1 assay (Roche, 1644807) 200 µL of the working reagent was added per slice and incubated for 45 min in the incubator. Negative controls were prepared by adding working reagent to slices that were incubated with 0.1% Saponin at 37°C for one hour. Each slice was measured in technical duplicates (75 µL per well in 96-well plate) and averaged.

### Cytokine secretion by MSD multiplex assay

Absolute levels of cytokines secreted by hPCLS were measured using MSD human chemokine panel (K15047D-2), human proinflammatory panel 1 (K15049D-2), IL-8 (K151ANB) and IP-10 kits (K151AVB-2) according to the manufacturer’s instructions. For chemokine and proinflammatory panels tissue culture supernatant was diluted 1:2, for IL-8 1:10, for IP-10 1:5.

### Cilia beating frequency (CBF) assay

The day following hPCLS preparation airway-containing hPCLS were manually selected and placed into MatTek dishes with 10-mm glass window (P35G-0-10-C) with 200 µL DMEM-BSA and covered with a cover slip to prevent slice from floating. Inverted Nikon-Ti-E microscope with environmental controlled box equipped with 20× 0.75NA air objective was used. Transmitted light images (512×512 px, 4×4 binning) were acquired at 250 fps for a total of 1000 frames along circumference of each airway. To examine temperature dependence of CBF, the same airway hPCLS were imaged at ambient temperature (t=26°C), at t=37°C and at 41°C. The effect of PI3Kδ inhibition on CBF was examined at t=37°C. CBF was quantified both manually and using automated Python-based pipeline. For manual analysis rectangular regions (15×3 µm) were selected along the airway border with beating cilia, converted to kymograms using ImageJ function Reslice and quantified using ImageJ function FFT (Fast Fourier transform). Custom-made Python-based script was developed to batch-analyze CBF movies (github link). In brief, for each pixel FFT was performed and highest power value and its corresponding frequency were noted. Pixels that had power values above given threshold (5–7 times above the median) were marked as positive for cilia beating and CBF values in the range of 2–30 Hz were extracted. More detailed documentation of the Python code functionality is provided on GitHub.

### Bead transport assay

Bead transport assay was performed simultaneously with CBF assay. Prior to covering hPCLS with a coverslip in a MatTek dish, 5,000 2-µm Nile Red FluoSpheres (ThermoFisher, F8825) dissolved in 2 µL DMEM-BSA were added. Bead transport was recorded on Nikon-Ti-E microscope equipped with 10× 0.45NA air objective. To increase number of tracked beads up to 10 time lapse images of the same airway (512×512 mm, 4×4 binning) were captured in 0.5 sec intervals for 30 sec in Cy3 channel. Beads were tracked using Manual Tracking ImageJ plugin. In addition to an average velocity of each bead, a maximum velocity was quantified by calculating a moving average of four consecutive steps for each bead, and taking the highest value using custom-made R script (available by request).

### Whole-mount hPCLS immunofluorescence and confocal imaging

hPCLS were fixed in 3% PFA in PBS for 24 h at 4°C, washed with PBS and blocked for non-specific antibody binding with 10% goat serum (Sigma, G9023) in PBS with 0.2% Triton X-100 (PBS-Tx) for 90 min at RT. Primary antibodies against Arl13b (Proteintech, 17711-1-AP, rabbit 1:500), p63 (Abcam, ab735, mouse 1:500), Muc5ac (Abcam, ab3649, mouse 1:500) were mixed in 1% goat serum in PBS-Tx and incubated for 22 h at RT. Following washing with PBS-Tx, hPCLS were incubated in a mix containing anti-rabbit Alexa488 (ThermoFisher, A-11008, 1:500), anti-mouse Alexa647 (ThermoFischer, A-21236, 1:500), Hoechst 33258 (Sigma, 861405, 0.4 µg/mL) and Phalloidin-Alexa568 (ThermoFisher, A12380, 1:600) in 1% goat serum in PBS-Tx for 2 h at RT. Washed hPCLS were embedded in mounting medium (ibidi, 50001) and sealed between two coverslips (24×50 mm, Carl Roth, LH25.1) using two-component silicon resin (Picodent, 1300-1000). Confocal images were acquired using Olympus FV3000 microscope using 60× 1.3NA silicon objective with effective xy resolution of 188 nm. Sub-Nyquist z-stacks (slice every 4–6 µm) were acquired along circumference of each airway from both sides to account for heterogeneity of the samples.

### Confocal image analysis

Automated 2D analysis of images was performed using custom-made ImageJ plugin (GitHub link). In brief, Phalloidin signal was used to define the airway border. Percentage of ciliated surface was measured along 15-px wide band along the airway epithelium border. Total number of cells on an image, number of cells in the airway lumen, basal cell number and mucus area were also quantified and normalized to an airway border length. For each quantified parameter median and coefficient of variation was quantified for each z-stack.

Nyquist z-stacks were reconstructed and analyzed using ImarisXT (Bitplane). Spot detection algorithm (estimated diameter 4.5 µm) was used to assign a spot to each nucleus of all cells (Hoechst channel) and separately to all basal cells (p63 channel). Resulting ratio was quantified and expressed as a percentage of basal cells. To quantify percent of ciliated airway surface in 3D, Imaris volume rendering algorithm was used for cilia and Phalloidin channels. Overlap of two volumes was quantified using Kiss and Run Imaris Extension.

### Statistical analysis

Statistical analysis was performed using GraphPad Prism 7. Statistical significance of differences was evaluated using unpaired *t*-test with Welch’s correction, paired *t*-test, one-way ANOVA or two-way ANOVA followed by Bonferroni post hoc test. Figure legends specified the statistical analysis used for the data in each panel. Differences were considered to be statistically significant when *P* < 0.05.

## Acknowledgements

Technical assistance from Christa Stolp from Biomaterial Bank Heidelberg (BMBH) in tissue assembling is gratefully acknowledged. We would also like to acknowledge the help of Augustin Amour for technical advice regarding the use GSK045.

## Supplementary Figures and tables

**Supplementary Figure S1.**
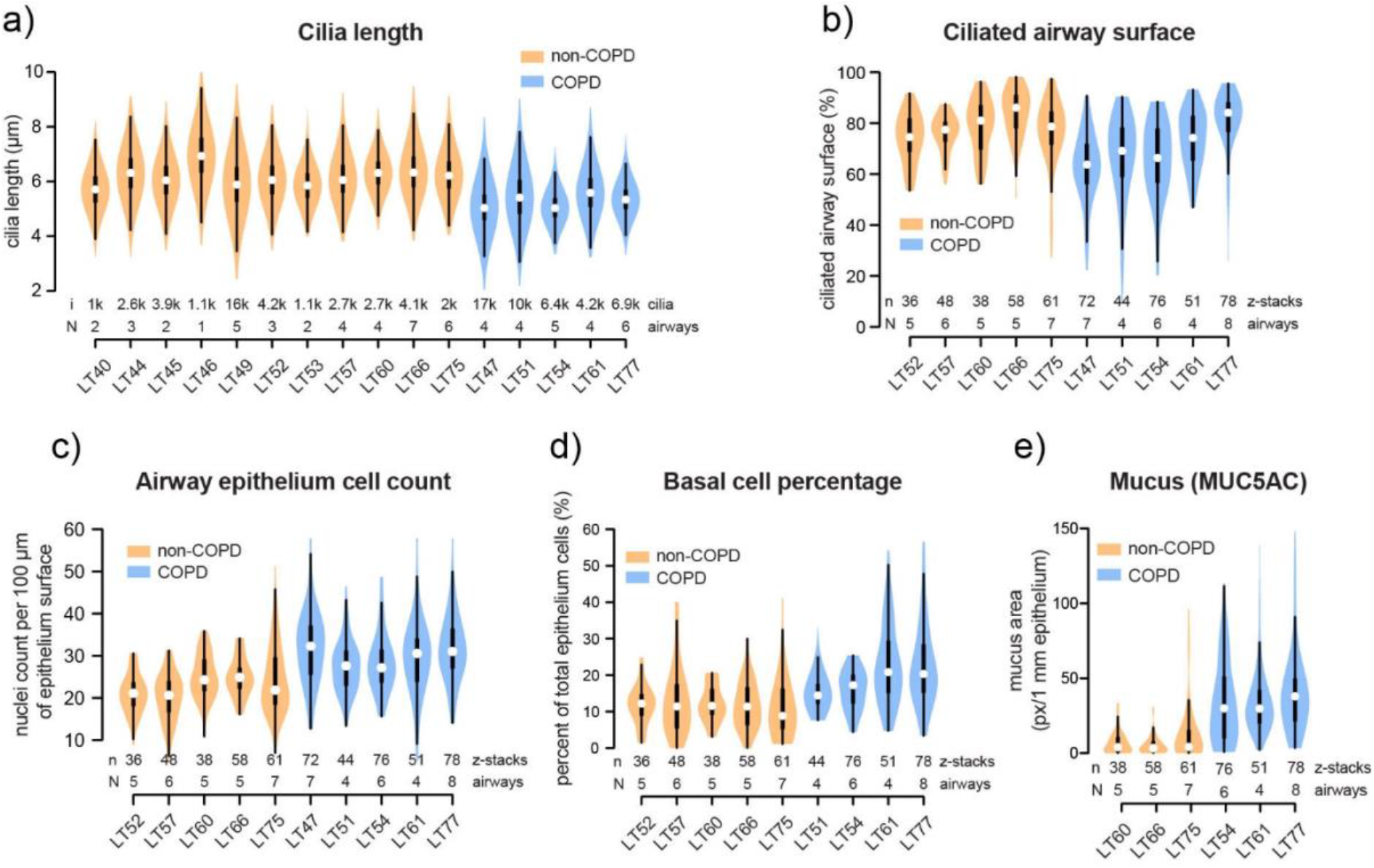
COPD-derived hPCLS retain disease-specific alterations. a–e) Measurements of cilia length (a), mucus (b), cililated airway surface (c), total airway epithelium cells (d) and percentage of basal cells in the airway epithelium (e) in the untreated non-COPD and COPD hPCLS. Panels show full distribution of a given parameter within each patient. White circles show the medians; box limits indicate the 25th and 75th percentiles; whiskers extend 1.5× the interquartile range from the 25th and 75th percentiles; polygons represent density estimates of data and extend to extreme values. N is the number of analyzed per donor, n is the total number of z-stacks, i is the number of manually measured cilia.

**Supplementary Figure S2.**
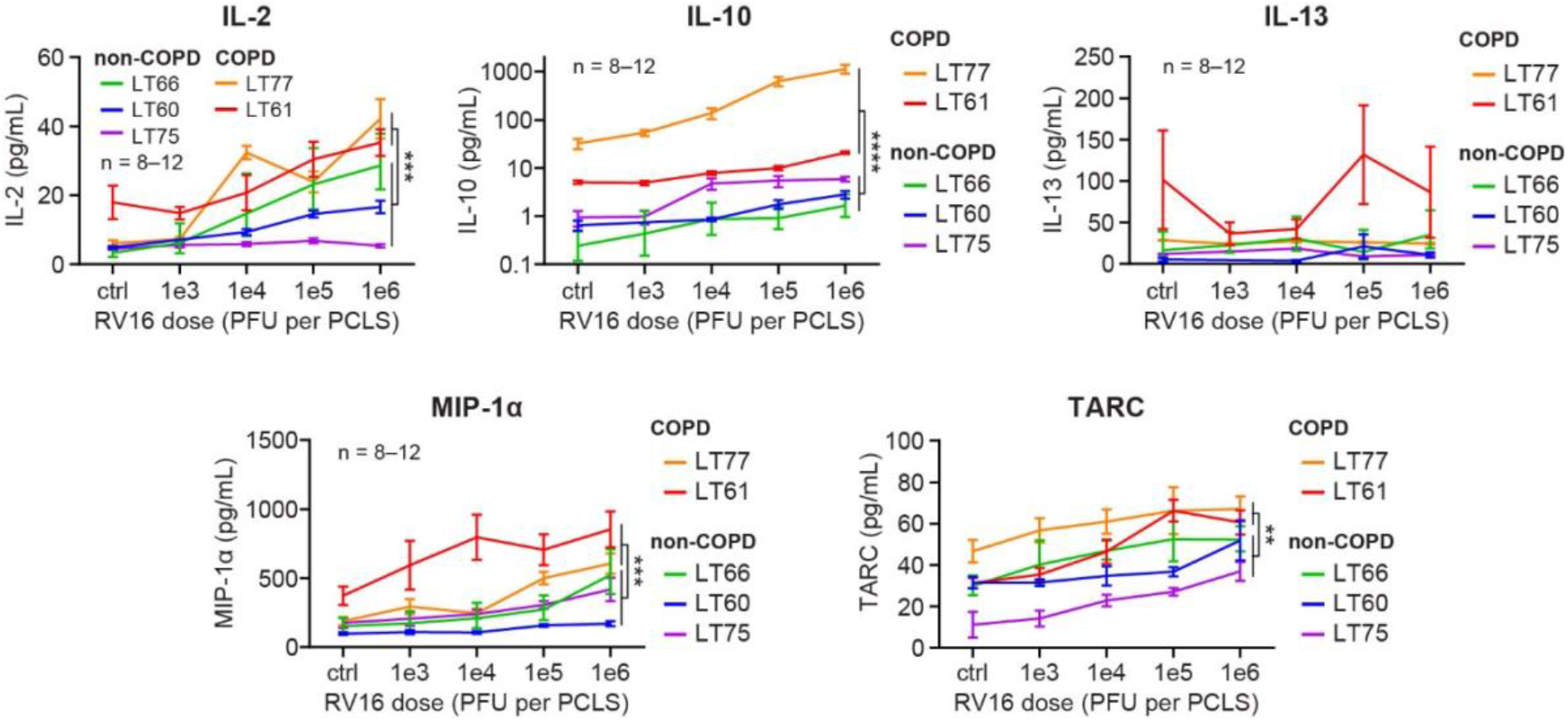
Rhinovirus infection induces stronger pro-inflammatory response in COPD hPCLS. hPCLS from three non-COPD and two COPD GOLD IV donors were infected with indicated amounts of rhinovirus 16 (RV16). Supernatants were collected 3 days after infection and analyzed for cytokine levels. Data correspond to mean and SEM, n is the number of hPCLS used per dose per donor. Statistical analysis by Factorial ANOVA with Bonferroni multiple comparison *post-hoc* test. **; ***; **** indicate significance with *P* < 0.01, 0.001; 0.0001. Full results of statistical analyses are shown in Supplementary table S2.

**Supplementary Figure S3.**
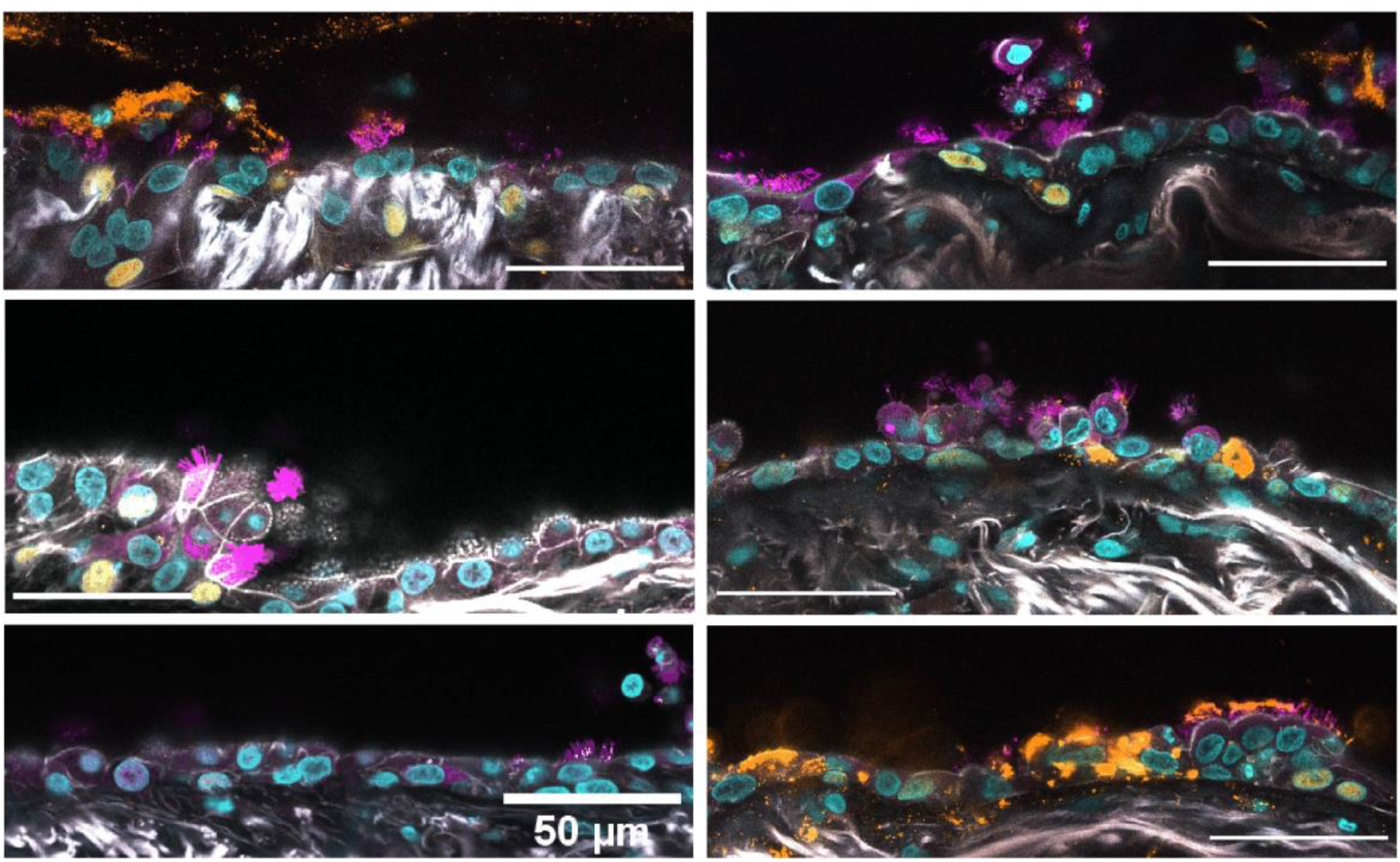
COPD small airway shedding after 1e6 PFU per hPCLS RV16 infection. COPD hPCLS were infected with 1e6 PFU per hPCLS of rhinovirus 16. Three days post infection, hPCLS were PFA-fixed, stained with corresponding antibodies and imaged on a confocal microscope. Scale bar 50 µm.

**Supplementary Figure S4.**
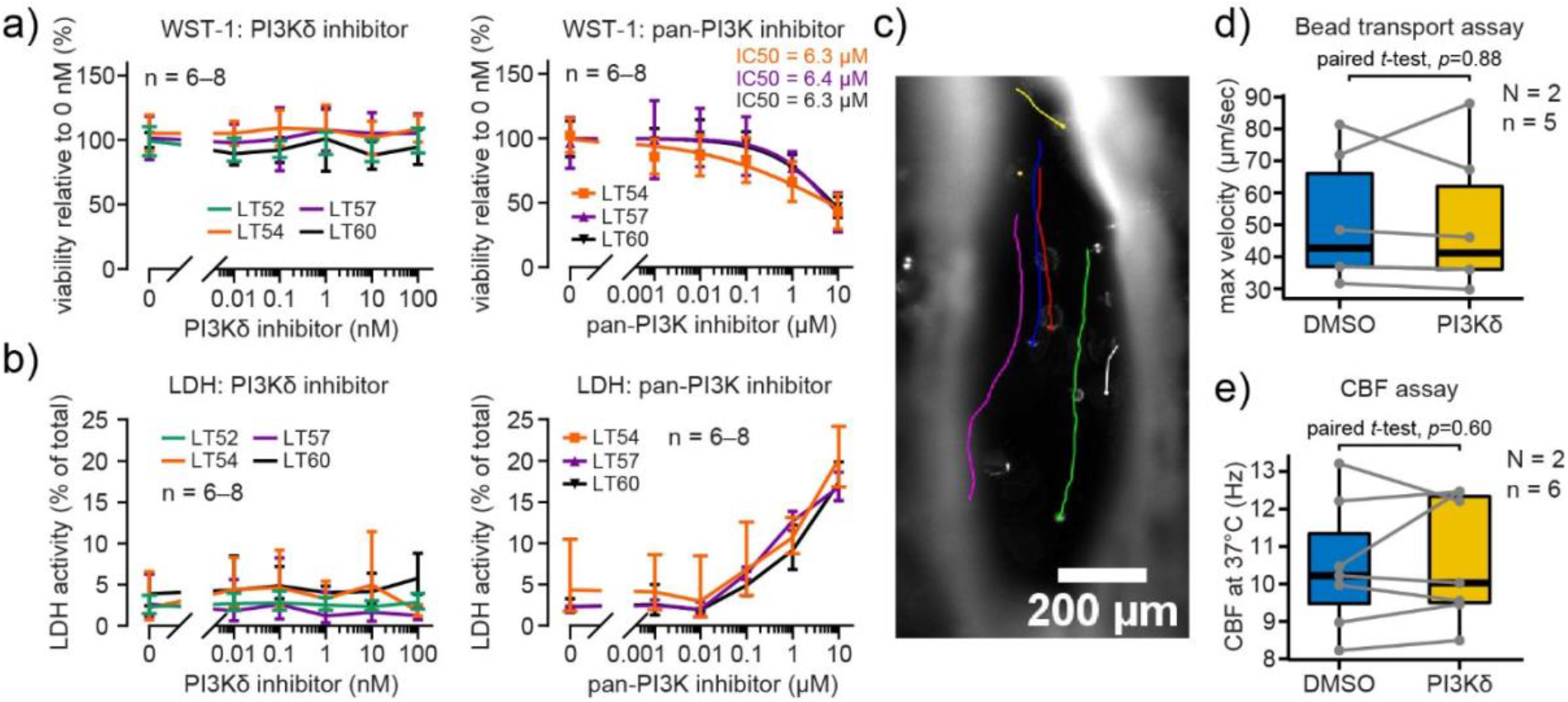
PI3Kδ inhibition has no apparent cytotoxicity in hPCLS. a) Evaluation of hPCLS mitochondrial activity using WST-1 viability assay. hPCLS were cultured for 3 days in increasing concentrations of PI3Kδ or pan-PI3K inhibitors. Data correspond to geometric mean and SD, n is the number of hPCLS used per condition per donor. b) Evaluation of LDH released from hPCLS after 3 days treatment with increasing concentrations of PI3Kδ or pan-PI3K inhibitors. Data correspond to mean and SEM, n is the number of hPCLS used per condition per donor. c) Representative image of the small airway and bead tracks. d, e) The effect of 5-hour treatment with 100 nM PI3Kδ inhibitor on bead transport velocity and CBF measured from the same hPCLS. Center lines show the medians; box limits indicate the 25th and 75th percentiles. N is the total number of donors, n is the total number of hPCLS.

**Supplementary Figure S5.**
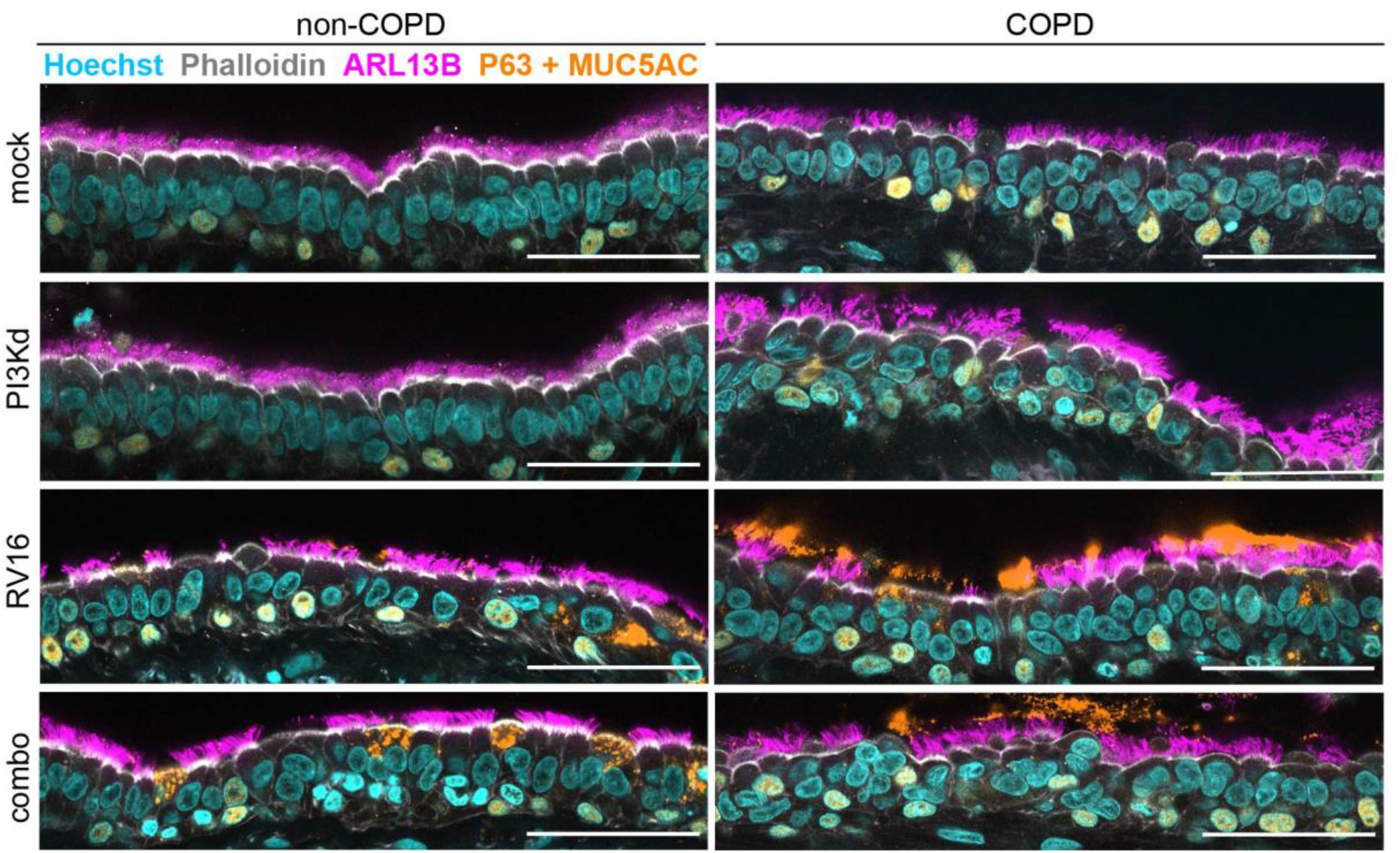
PI3Kδ inhibition differentially impacts integrity of COPD small airway epithelium after rhinovirus infection. hPCLS from three non-COPD and two COPD GOLD IV donors were left untreated (mock), or treated with 100 nM PI3Kδ inhibitor (PI3Kd), or infected with 1e5 PFU/hPCLS of rhinovirus 16 (RV16), or first pre-treated with PI3Kδ for five hours and then infected with RV16 (combo). Three days post treatment hPCLS were PFA-fixed, stained with corresponding antibodies and imaged on a confocal microscope. Representative images for each condition of non-COPD and COPD airways are shown in addition to images shown in Figure 5. Scale bar 50 µm.

**Supplementary Figure S6.**
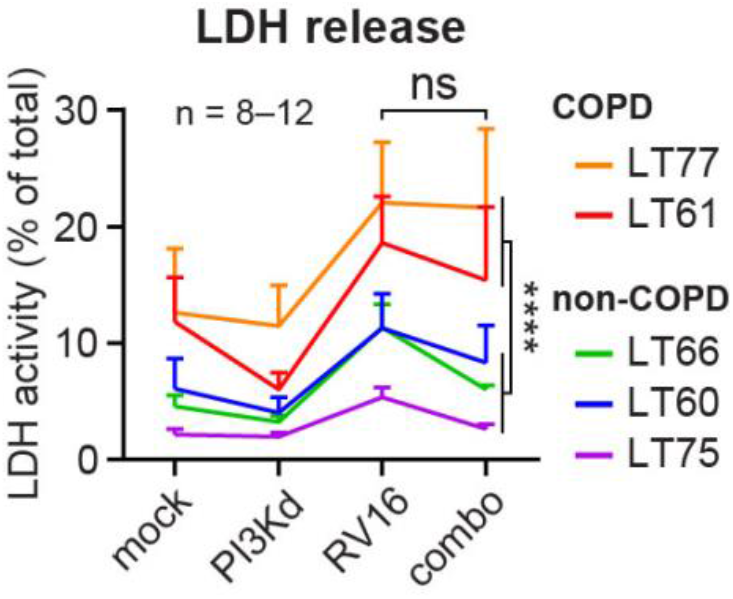
PI3Kδ inhibition does not alter rhinovirus-induced LDH release by non-COPD and COPD hPCLS. hPCLS from three non-COPD and two COPD GOLD IV donors were left untreated (mock), or treated with 100 nM PI3Kδ inhibitor (PI3Kd), or infected with 1e5 PFU/hPCLS of rhinovirus 16 (RV16), or first pre-treated with PI3Kδ for five hours and then infected with RV16 (combo). Three days post infection supernatants were collected and analyzed for LDH activity. Data correspond to mean and SEM, n is the number of hPCLS used per dose per donor. Statistical analysis by Factorial ANOVA with Bonferroni multiple comparison *post-hoc* test. **** indicate significance with 0.000, ns: not significant. Full results of statistical analyses are shown in supplementary table S4.

**Supplementary Table S1.**
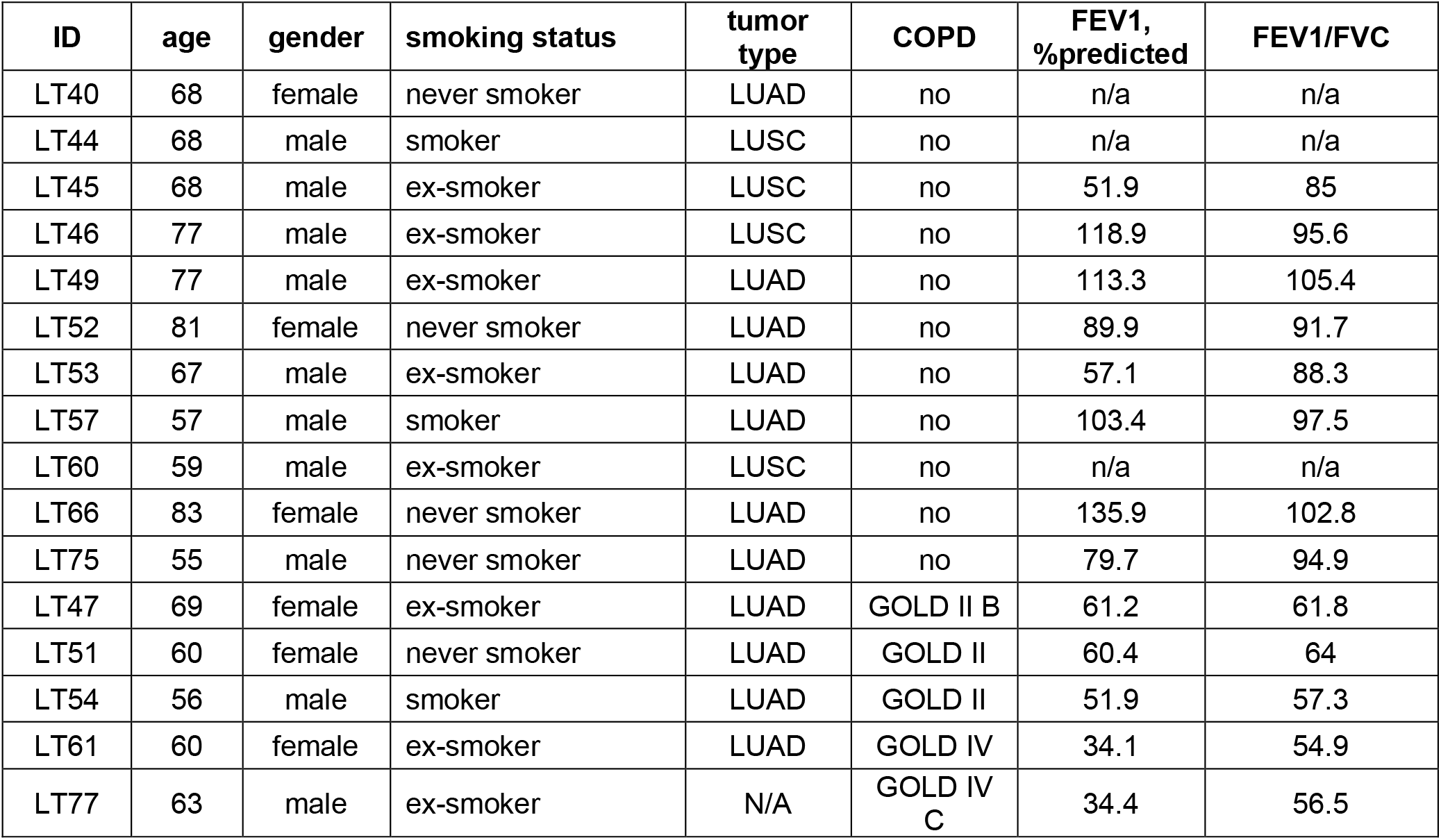
Clinical characteristics of patients included in this study

| ID | age | gender | smoking status | tumor type | COPD | FEV1, %predicted | FEV1/FVC |
| --- | --- | --- | --- | --- | --- | --- | --- |
| LT40 | 68 | female | never smoker | LUAD | no | n/a | n/a |
| LT44 | 68 | male | smoker | LUSC | no | n/a | n/a |
| LT45 | 68 | male | ex-smoker | LUSC | no | 51.9 | 85 |
| LT46 | 77 | male | ex-smoker | LUSC | no | 118.9 | 95.6 |
| LT49 | 77 | male | ex-smoker | LUAD | no | 113.3 | 105.4 |
| LT52 | 81 | female | never smoker | LUAD | no | 89.9 | 91.7 |
| LT53 | 67 | male | ex-smoker | LUAD | no | 57.1 | 88.3 |
| LT57 | 57 | male | smoker | LUAD | no | 103.4 | 97.5 |
| LT60 | 59 | male | ex-smoker | LUSC | no | n/a | n/a |
| LT66 | 83 | female | never smoker | LUAD | no | 135.9 | 102.8 |
| LT75 | 55 | male | never smoker | LUAD | no | 79.7 | 94.9 |
| LT47 | 69 | female | ex-smoker | LUAD | GOLD II B | 61.2 | 61.8 |
| LT51 | 60 | female | never smoker | LUAD | GOLD II | 60.4 | 64 |
| LT54 | 56 | male | smoker | LUAD | GOLD II | 51.9 | 57.3 |
| LT61 | 60 | female | ex-smoker | LUAD | GOLD IV | 34.1 | 54.9 |
| LT77 | 63 | male | ex-smoker | N/A | GOLD IV C | 34.4 | 56.5 |

## REFERENCE

1. Lozano, R. et al. Global and regional mortality from 235 causes of death for 20 age groups in 1990 and 2010: A systematic analysis for the Global Burden of Disease Study 2010. Lancet 380, (2012).

2. Soler-Cataluña, J. J. et al. Severe acute exacerbations and mortality in patients with chronic obstructive pulmonary disease. Thorax 60, (2005).

3. Wilkinson, T. M. A. et al. A prospective, observational cohort study of the seasonal dynamics of airway pathogens in the aetiology of exacerbations in COPD. Thorax 72, (2017).

4. Mallia, P. et al. Experimental rhinovirus infection as a human model of chronic obstructive pulmonary disease exacerbation. Am. J. Respir. Crit. Care Med. 183, (2011).

5. Lipson, D. A. et al. FULFIL Trial: Once-daily triple therapy for patients with chronic obstructive pulmonary disease. Am. J. Respir. Crit. Care Med. 196, (2017).

6. Knight, D. A. & Holgate, S. T. The airway epithelium: structural and functional properties in health and disease. Respirology 8, 432–446 (2003).

7. Crystal, R. G., Randell, S. H., Engelhardt, J. F., Voynow, J. & Sunday, M. E. Airway epithelial cells: current concepts and challenges. Proc. Am. Thorac. Soc. 5, 772–777 (2008).

8. Papadopoulos, N. G. et al. Rhinoviruses infect the lower airways. J. Infect. Dis. 181, (2000).

9. Holgate, S. T. Epithelial damage and response. Clin. Exp. allergy J. Br. Soc. Allergy Clin. Immunol. 30 Suppl 1, 37–41 (2000).

10. Puchelle, E., Zahm, J.-M., Tournier, J.-M. & Coraux, C. Airway epithelial repair, regeneration, and remodeling after injury in chronic obstructive pulmonary disease. Proc. Am. Thorac. Soc. 3, 726–733 (2006).

11. Vanhaesebroeck, B. et al. P110delta, a novel phosphoinositide 3-kinase in leukocytes. Proc. Natl. Acad. Sci. U. S. A. 94, 4330–4335 (1997).

12. To, Y. et al. Targeting phosphoinositide-3-kinase-δ with theophylline reverses corticosteroid insensitivity in chronic obstructive pulmonary disease. Am. J. Respir. Crit. Care Med. 182, (2010).

13. Sapey, E. et al. Behavioral and structural differences in migrating peripheral neutrophils from patients with chronic obstructive pulmonary disease. Am. J. Respir. Crit. Care Med. 183, (2011).

14. Milara, J. et al. Roflumilast N-oxide reverses corticosteroid resistance in neutrophils from patients with chronic obstructive pulmonary disease. J. Allergy Clin. Immunol. 134, (2014).

15. Lucas, C. L. et al. Dominant-activating germline mutations in the gene encoding the PI(3)K catalytic subunit p110δ result in T cell senescence and human immunodeficiency. Nat. Immunol. 15, 88–97 (2014).

16. Angulo, I. et al. Phosphoinositide 3-kinase δ gene mutation predisposes to respiratory infection and airway damage. Science 342, 866–871 (2013).

17. Stark, A.-K. et al. PI3Kδ hyper-activation promotes development of B cells that exacerbate Streptococcus pneumoniae infection in an antibody-independent manner. Nat. Commun. 9, 3174 (2018).

18. Churg, A., Sin, D. D. & Wright, J. L. Everything prevents emphysema: are animal models of cigarette smoke-induced chronic obstructive pulmonary disease any use? Am. J. Respir. Cell Mol. Biol. 45, 1111–1115 (2011).

19. Mestas, J. & Hughes, C. C. W. Of mice and not men: differences between mouse and human immunology. J. Immunol. 172, 2731–2738 (2004).

20. Fehrenbach, H. Animal models of chronic obstructive pulmonary disease: some critical remarks. Pathobiology 70, 277–283

21. Sanderson, M. J. Exploring lung physiology in health and disease with lung slices. Pulm. Pharmacol. Ther. 24, 452–465 (2011).

22. Khan, M. M. et al. An integrated multiomic and quantitative label-free microscopy-based approach to study pro-fibrotic signalling in ex vivo human precision-cut lung slices. Eur. Respir. J. 2000221 (2020). doi:10.1183/13993003.00221-2020

23. Clary-Meinesz, C. F., Cosson, J., Huitorel, P. & Blaive, B. Temperature effect on the ciliary beat frequency of human nasal and tracheal ciliated cells. Biol. cell 76, 335–338 (1992).

24. Ramshaw, I. A. et al. Cytokines and immunity to viral infections. Immunol. Rev. 159, 119–135 (1997).

25. Down, K. et al. Optimization of Novel Indazoles as Highly Potent and Selective Inhibitors of Phosphoinositide 3-Kinase δ for the Treatment of Respiratory Disease. J. Med. Chem. 58, 7381–7399 (2015).

26. Footitt, J. et al. Oxidative and Nitrosative Stress and Histone Deacetylase-2 Activity in Exacerbations of COPD. Chest 149, 62–73 (2016).

27. Zou, Y. et al. Serum IL-1β and IL-17 levels in patients with COPD: associations with clinical parameters. Int. J. Chron. Obstruct. Pulmon. Dis. 12, 1247–1254 (2017).

28. Schneider, D. et al. Increased cytokine response of rhinovirus-infected airway epithelial cells in chronic obstructive pulmonary disease. Am. J. Respir. Crit. Care Med. 182, 332–340 (2010).

29. Baines, K. J. et al. Novel immune genes associated with excessive inflammatory and antiviral responses to rhinovirus in COPD. Respir. Res. 14, 15 (2013).

30. McKendry, R. T. et al. Dysregulation of Antiviral Function of CD8(+) T Cells in the Chronic Obstructive Pulmonary Disease Lung. Role of the PD-1-PD-L1 Axis. Am. J. Respir. Crit. Care Med. 193, 642–651 (2016).

31. Singanayagam, A. et al. A short-term mouse model that reproduces the immunopathological features of rhinovirus-induced exacerbation of COPD. Clin. Sci. (Lond). 129, 245–258 (2015).

32. Ganesan, S. et al. Aberrantly activated EGFR contributes to enhanced IL-8 expression in COPD airways epithelial cells via regulation of nuclear FoxO3A. Thorax 68, 131–141 (2013).

33. Hsu, A. C.-Y. et al. Targeting PI3K-p110α Suppresses Influenza Virus Infection in Chronic Obstructive Pulmonary Disease. Am. J. Respir. Crit. Care Med. 191, 1012–1023 (2015).

34. Sajjan, U. et al. Elastase- and LPS-exposed mice display altered responses to rhinovirus infection. Am. J. Physiol. Lung Cell. Mol. Physiol. 297, L931–44 (2009).

35. Almansa, R. et al. Host response cytokine signatures in viral and nonviral acute exacerbations of chronic obstructive pulmonary disease. J. Interf. cytokine Res. Off. J. Int. Soc. Interf. Cytokine Res. 31, 409–413 (2011).

36. Robbins, C. S. et al. Cigarette smoke impacts immune inflammatory responses to influenza in mice. Am. J. Respir. Crit. Care Med. 174, 1342–1351 (2006).

37. Arruda, E. et al. Localization of human rhinovirus replication in the upper respiratory tract by in situ hybridization. J. Infect. Dis. 171, 1329–1333 (1995).

38. Bardin, P. G. et al. Amplified rhinovirus colds in atopic subjects. Clin. Exp. allergy J. Br. Soc. Allergy Clin. Immunol. 24, 457–464 (1994).

39. Mosser, A. G. et al. Similar frequency of rhinovirus-infectible cells in upper and lower airway epithelium. J. Infect. Dis. 185, 734–743 (2002).

40. Cakebread, J. A. et al. Rhinovirus-16 induced release of IP-10 and IL-8 is augmented by Th2 cytokines in a pediatric bronchial epithelial cell model. PLoS One 9, e94010 (2014).

41. Marwick, J. A. et al. A role for phosphoinositol 3-kinase delta in the impairment of glucocorticoid responsiveness in patients with chronic obstructive pulmonary disease. J. Allergy Clin. Immunol. 125, 1146–1153 (2010).

42. Cai, Y. et al. Immunological characterization of HM5023507, an orally active PI3Kδ/γ inhibitor. Pharmacol. Res. Perspect. 8, e00559 (2020).

43. Khindri, S. et al. A Multicentre, Randomized, Double-Blind, Placebo-Controlled, Crossover Study To Investigate the Efficacy, Safety, Tolerability, and Pharmacokinetics of Repeat Doses of Inhaled Nemiralisib in Adults with Persistent, Uncontrolled Asthma. J. Pharmacol. Exp. Ther. 367, 405–413 (2018).

44. Cahn, A. et al. Safety, pharmacokinetics and dose-response characteristics of GSK2269557, an inhaled PI3Kδ inhibitor under development for the treatment of COPD. Pulm. Pharmacol. Ther. 46, 69–77 (2017).

45. Begg, M. et al. Exploring PI3Kδ Molecular Pathways in Stable COPD and Following an Acute Exacerbation, Two Randomized Controlled Trials. Int. J. Chron. Obstruct. Pulmon. Dis. 16, 1621–1636 (2021).

46. Chang, S. C. Microscopic properties of whole mounts and sections of human bronchial epithelium of smokers and nonsmokers. Cancer 10, 1246–1262 (1957).

47. Leopold, P. L. et al. Smoking is associated with shortened airway cilia. PLoS One 4, e8157 (2009).

48. Hessel, J. et al. Intraflagellar transport gene expression associated with short cilia in smoking and COPD. PLoS One 9, e85453 (2014).

49. Boers, J. E., Ambergen, A. W. & Thunnissen, F. B. Number and proliferation of basal and parabasal cells in normal human airway epithelium. Am. J. Respir. Crit. Care Med. 157, 2000–2006 (1998).

50. Hogg, J. C. & Timens, W. The pathology of chronic obstructive pulmonary disease. Annu. Rev. Pathol. 4, 435–459 (2009).

51. Chung, K. F. & Adcock, I. M. Multifaceted mechanisms in COPD: inflammation, immunity, and tissue repair and destruction. Eur. Respir. J. 31, 1334–1356 (2008).

52. Sadhu, C., Dick, K., Tino, W. T. & Staunton, D. E. Selective role of PI3K delta in neutrophil inflammatory responses. Biochem. Biophys. Res. Commun. 308, 764–769 (2003).

53. Condliffe, A. M. et al. Sequential activation of class IB and class IA PI3K is important for the primed respiratory burst of human but not murine neutrophils. Blood 106, 1432–1440 (2005).

54. Poolman, T. M., Ng, L. L., Farmer, P. B. & Manson, M. M. Inhibition of the respiratory burst by resveratrol in human monocytes: correlation with inhibition of PI3K signaling. Free Radic. Biol. Med. 39, 118–132 (2005).

55. Ali, K. et al. Essential role for the p110delta phosphoinositide 3-kinase in the allergic response. Nature 431, 1007–1011 (2004).

56. Hogg, J. C. et al. The nature of small-airway obstruction in chronic obstructive pulmonary disease. N. Engl. J. Med. 350, 2645–2653 (2004).

57. Kosanovic, D. et al. Histological characterization of mast cell chymase in patients with pulmonary hypertension and chronic obstructive pulmonary disease. Pulm. Circ. 4, 128–136 (2014).

58. Keatings, V. M. & Barnes, P. J. Granulocyte activation markers in induced sputum: comparison between chronic obstructive pulmonary disease, asthma, and normal subjects. Am. J. Respir. Crit. Care Med. 155, 449–453 (1997).

59. Rohde, G. et al. Inflammatory response in acute viral exacerbations of COPD. Infection 36, 427–433 (2008).

